# Adipocyte-Specific Ablation Of PU.1 Promotes Energy Expenditure and Ameliorates Metabolic Syndrome In Aging Mice

**DOI:** 10.1101/2021.08.29.458103

**Authors:** Keyun Chen, Alejandra De Angulo, Xin Guo, Aditya More, Scott A. Ochsner, Eduardo Lopez, David Saul, Weijun Pang, Yuxiang Sun, Neil J. McKenna, Qiang Tong

**Affiliations:** USDA/ARS Children’s Nutrition Research Center, Department of Pediatrics, Baylor College of Medicine, Houston, Texas, USA; Department of Nutrition and Food Hygiene, School of Public Health, Cheeloo College of Medicine, Shandong University, Jinan, China; Department of Molecular and Cellular Biology, Baylor College of Medicine, Houston, Texas, USA; Northwestern University of Agriculture and Forestry, Yangling, Shaanxi, China; Department of Nutrition, Texas A&M University, College Station, Texas, USA; Department of Molecular Physiology & Biophysics, Department of Medicine, Baylor College of Medicine, Huffington Center on Aging, Houston, Texas, USA

**Keywords:** Spi1/PU.1, Adipocyte, Aging, Transcription, Obesity, Insulin Resistance, Inflammation, Energy Expenditure, Thermogenesis

## Abstract

**Objective:** Although PU.1/Spi1 is known as a master regulator for macrophage development and function, we have reported previously that it is also expressed in adipocytes and is transcriptionally induced in obesity. Here, we investigated the role of adipocyte PU.1 in the development of age-associated metabolic syndrome.

**Methods:** We generated mice with adipocyte specific PU.1 knockout, assessed metabolic changes in young and aged PU.1^fl/fl^ (control) and AdipoqCre PU.1^fl/fl^(aPU.1KO) mice, including body weight, body composition, energy expenditure and glucose homeostasis. We also performed transcriptional analyses using RNA-Sequencing of adipocytes from these mice.

**Results:** aPU.1KO mice have elevated energy expenditure at a young age and decreased adiposity and increased insulin sensitivity in later life. Corroborating these observations, transcriptional network analysis indicated the existence of validated, aPU.1-modulated regulatory hubs that direct inflammatory and thermogenic gene expression programs.

**Conclusions:** Our data provide evidence for a previously uncharacterized role of PU.1 in the development of age-associated obesity and insulin resistance.

## 1. INTRODUCTION

Systemic insulin resistance is a major global public health concern. Although it impacts numerous metabolic organs including skeletal muscle, liver and adipose tissue, evidence indicates that adipose tissue is one of the primary origins of systemic insulin resistance. It is well documented, for example, that obesity induces chronic low-grade inflammation in adipose tissue, a key event leading to systemic insulin resistance and metabolic syndrome ^1^. During obesity, adipose tissue secretes elevated levels of pro-inflammatory cytokines, such as TNF-α, IL-1β, IL-6 and MCP-1^2^. Obesity-associated adipose inflammation, for instance, is characterized by increased infiltration of macrophages ^3, 4^ and other immune cells ^5–8^, and macrophage infiltration has been shown to be stimulated by obese adipose tissue expression of Ccl2/MCP-1 ^9, 10^. Although the role of macrophages is well-established in the inflammatory processes accompanying insulin resistance in adipose tissue, accumulating evidence implicates adipocytes as active participants in these processes.

PU.1 (encoded by *Spi1*) is a member of the ETS family of transcription factors ^11^ with historically well-characterized roles in the development of myeloid and lymphoid lineages, in particular macrophages and granulocytes ^12–14^. Functions for PU1.1 have also been established in lineage establishment of microglia ^15^, dendritic cells ^16^, and osteoclasts ^17^. Additionally, numerous lines of evidence cast macrophage PU.1 in a central role in the coordination of inflammatory transcriptional programs in macrophages. For example, macrophages lacking PU.1 are deficient in lipopolysaccharide (LPS) induction of *Tlr4*, *Ptgs2* (encoding COX-2), *Tnf*, *Il1b*, *Il6*, and *Ccl2/MCP-1* ^18^. In contrast to the volume of studies on PU.1 function in macrophages however, relatively little is known about its role in adipocytes. We previously reported that PU.1 was expressed in adipocytes, and that adipose tissue PU.1 expression was greatly increased in mouse models of obesity ^19^. Consistent with its pro-inflammatory role in macrophages, we found that depletion of PU.1 in cultured adipocytes led to decreased reactive oxygen species (ROS) production, increased insulin sensitivity, and reduced expression of signature obese adipose tissue cytokines, including TNF-α, IL-1β, and IL-6 ^20^. In addition, Lackey and colleagues have recently reported that mice with adipocyte-specific PU.1 knockout showed improved insulin sensitivity on a high fat diet, without any difference in body weight ^21^.

The age-associated metabolic syndrome may have characteristics different from that caused by diet-induced obesity ^22^. Here, we investigated the adipocyte-specific functions of PU.1 in mice, particularly during aging. We found that male mice with adipocyte-specific ablation of PU.1 had elevated energy expenditure, and were protected against age-associated obesity, with increased insulin sensitivity and increased glucose tolerance. Mechanistically, we performed validated informatic analyses that connect PU.1-modulated transcriptional hubs to the observed physiological changes.

## 2. EXPERIMENTAL PROCEDURES

### 2.1 Animals Experiments

PU.1^fl/+^ mice, containing loxP sites flanking exon 5 of the *Spi1* gene, were obtained from Dr. Stephen Nutts ^23^. These mice are on C57BL/6 genetic background and were used successfully for tissue specific knockout (59). Mice with adipocyte-specific knockout of PU.1 (aPU.1KO) were generated by crossing Adiponectin-Cre mice (Adipoq-Cre, The Jackson Laboratory) ^24^ with PU.1 floxed mice. Mice were fed on a standard chow diet (Lab Diet 5053; Purina Mills). All procedures used in animal experiments were approved by the Institution of Animal Care and Use Committee at Baylor College of Medicine. Mice were maintained under conditions of controlled temperature (∼75°F) and illumination (12-hour light/12-hour dark cycle, 6am to 6pm) with free access to water.

### 2.2 Glucose Tolerance Test (GTT) and Insulin Tolerance Test (ITT)

Mice were fasted for overnight (GTT) or 4 h (ITT) and received an intraperitoneal injection of pyruvate (2g/kg), D-glucose (2 g/kg) or insulin (1 IU/kg). Blood glucose was measured by glucose meter (TrueTest Glucose Meter and Strips) from the tail vein before and at 15, 30, 60, and 120 min after the bolus pyruvate, glucose or insulin injection.

### 2.3 Body Composition Measurement and Calorimetry Experiment

Mice body composition was measured using the EchoMRI-100™ quantitative NMR instrument (Echo Medical Systems).

### 2.4 Indirect calorimetry

Indirect calorimetry was measured using a computer-controlled, open-circuit system (Oxymax System) as part of an integrated Comprehensive Lab Animal Monitoring System (CLAMS; Columbus Instruments, Columbus, OH, USA). Mice were singly housed in individual cages in adaptation for 3 days, followed by measurement for 4 days. On the last day, food was removed, and mice were fasted for 6 hours. Oxygen consumption (VO2) and carbon dioxide production (VCO2) were measured for each chamber and calculated by Oxymax software (v. 5.9). Energy expenditure was calculated as EE = 3.815 × VO2 + 1.232 × VCO2. The basal energy expenditure was calculated based on the three lowest EE time points during the fasting period.

### 2.5 Adipose tissue fractionation

Epididymal fat pads from mice were minced in Krebs-Ringer phosphate buffer and digested with 1 mg/ml collagenase type I (Worthington Biochemical) at 37°C for 1 h as described in the literature (58). Digested tissue was filtered through a nylon mesh and centrifuged at 500 rpm for 10 min. The top layer (adipocyte fraction) was collected. The remaining was centrifuged again at 1,500 rpm for 10 min, and the pellet (stromal-vascular fraction) was collected. Proteins were extracted from both fractions for Western blot analysis.

### 2.6 RNA-sequencing

Epididymal adipocytes RNA were extracted using the RNeasy Lipid Tissue Mini Kit (QIAGEN). The cells were homogenized in 1 ml QIAzol lysis reagent and centrifuged at 12,000 g for 10 min at 4°C. The lysates under the lipid layer were transferred to a fresh tube and extracted with 200 μl chloroform, centrifuged at 12,000 g for 15 min at 4°C. The upper aqueous phase was transferred out and mixed with 1 volume of 70% ethanol. The samples were then applied to RNeasy Mini spin column and centrifuged at room temperature for 15 s at 8,000 g. The columns were washed once with 700 μl Buffer RW1, and twice with 500 μl Buffer RPE. The RNA samples were eluded with 30–50 μl RNase-free water. RNA-sequencing was performed by Novogene Corporation Inc. (Sacramento, CA).

### 2.7 RNA-Seq Analysis

Sequencing was performed on adipocyte samples from PU.1 fl/fl (control) and PU.1 fl/fl-AdipoqCre (PU.1 knockout) mice, with 3 replicates in each group. Sequencing reads were quantified using Salmon with the option-validateMappings for a more sensitive mapping scheme^25^. Transcript-level counts were summed to the gene-level for differential expression analysis using DESeq2 ^26^.

### 2.8 Real-time PCR

Adipocytes were isolated from mice gonadal adipose tissue. Total RNA of adipocytes was isolated using TRIzol Reagent (Invitrogen, Carlsbad, CA) following the manufacturer’s instructions. The cDNA was synthesized using the SuperScript III First-Strand Synthesis System for RT-PCR (Invitrogen, Carlsbad, CA). qRT-PCR reactions were performed using iTaq Universal SYBR Green in a CFX96 Touch Real-Time PCR Detection System (Bio-Rad). The ΔCt method (2–ΔCt) was used to calculate the relative mRNA expression level of each gene. Specific gene expression was normalized to 18S ribosomal RNA. Sequences of primers used for real-time PCR were as follows: MCP-1-F 5’-GAAGGAATGGGTCCAGACAT-3’ and MCP-1-R 5’-ACGGGTCAACTTCACATTCA-3’; TNFα-F 5’-ACGGGTCAACTTCACATTCA-3’ and TNFα-R 5′-CTGATGAGAGGGAGGCCATT-3′; UCP-1-R 5’- AGCCACCACAGAAAGCTTGTCAAC-3’ and UCP-1-R 5’- ACAGCTTGGTACGCTTGGGTACTG-3’; PGC1α-F 5’- GTCAACAGCAAAAGCCACAA-3’ and PCG1α-R 5’-TCTGGGGTCAGAGGAAGAGA-3’; ChREBP-F 5’- AGTGCTTGAGCCTGGCCTAC-3’ and ChREBP-R 5’-TTGTTCAGGCGGATCTTGTC-3’; ′; 18S ribosomal RNA-F 5′-AACGAGACTCTGGCATGCTAACTAG-3′ and 18S ribosomal RNA-R 5′- CGCCACTTGTCCCTCTAAGAA-3′. The expression levels of genes of interest were normalized by the levels of 18S RNA.

### 2.9 Western blot analysis

Cells were lysed in lysis buffer (50 mM Tris, 50 mM KCl, 20 mM NaF, 1 mM Na_3_VO_4_, 10 mM EDTA, 1% NP-40, 1 mM PMSF, 5 μg/ml leupeptin, pH 8.0). Protein concentration was determined with BCA protein assay kit (Pierce, Rockford, IL). Twenty microgram proteins of each sample were separated by SDS-PAGE and electro-transferred to nitrocellulose membrane for immunoblot analysis. The following antibodies were used: anti-PU.1 (Santa Cruz Biotechnology, Santa Cruz, CA; sc-352, 1:500), anti-a-tubulin (Sigma, St. Louis, MO; T5168, 1:100,000), HRP-conjugated anti-mouse (Bio-Rad, Richmond, CA; 170-6516, 1:30,000), anti-rabbit (Bio-Rad, 170-6515, 1:30,000. The SuperSignal West Pico Chemiluminescent kit (Pierce, Rockford, IL) was used as substrates.

### 2.10 3T3-L1 adipogenesis consensome

Full details of the methods and principles underlying consensome analysis can be found in the original publication ^27^. Briefly, five transcriptomic datasets (GSE2192, GSE60745, GSE14004, GSE12929, GSE20696) generated from 3T3-L1 cells treated with a standard adipogenic cocktail were organized into appropriate contrasts comparing gene expression levels at different time points to day 0 expression levels. These contrasts were then processed by the consensome pipeline implemented in R as previously described. For each transcript, the algorithm counts the number of experiments where the significance for differential expression is <0.05, then generates the binomial probability, referred to as the consensome p-value (CPV), of observing that many or more nominally significant experiments out of the number of experiments in which the transcript was assayed, given a true probability of 0.05. Genes were ranked firstly by CPV, then by geometric mean fold change (GMFC). The 3T3-L1 adipogenesis consensome was validated against the GSEA adipogenesis Hallmark gene set using a hypergeometric test implemented in R as described in the results.

### 2.11 High confidence transcriptional target intersection analysis

Node and node family consensomes are gene lists ranked according to measures of the strength of their regulatory relationship with upstream signaling pathway nodes derived from independent publicly archived transcriptomic or ChIP-Seq datasets. In the case of ChIP-Seq datasets, the strength of the regulatory relationship is inferred from the mean ChIP-Atlas ^28^ MACS2 peak strength across available archived ChIP-Seq datasets in which a given pathway node is the IP antigen. In the case of transcriptomic datasets, the strength of the regulatory relationship is inferred from the frequency of significant differential expression of a given gene across independent experiments involving perturbation of a member of a given node family ^27^. Genes in the 95th percentile of a given node consensome were designated high confidence transcriptional targets (HCTs) for that node and used as the input for the HCT intersection analysis using the Bioconductor GeneOverlap analysis package implemented in R as previously described ^29^. For both consensome and HCT intersection analysis, *P* values were adjusted for multiple testing by using the method of Benjamini & Hochberg to control the false discovery rate as implemented with the p.adjust function in R, to generate *Q* values. Evidence for a transcriptional regulatory relationship between a node and a gene set was inferred from a larger intersection between the gene set and HCTs for a given node or node family than would be expected by chance after FDR correction (Q < 0.05).

### 2.12 Mammalian Phenotype Ontology analysis

Genes mapping to MPO terms phenotypes were retrieved from MGI ^30^. A hypergeometric test implemented in GraphPad Prism 7.0 was used to estimate the over-representation, relative to their distribution in all 691 nodes, of nodes encoded by MPO term-mapped genes among nodes with significant HCT intersections with aPU.1KO-induced or -repressed genes.

### 2.13 Statistics

The data are represented as the mean ± standard error. Statistical significance was determined using the two-tail Student’s *t*-test. *P* < 0.05 was considered to be statistically significant.

## 3. RESULTS

### 3.1 Generation of adipocyte-specific PU.1 knockout mice

To investigate the systemic functions of aPU.1 *in vivo*, we generated mice with adipose-specific knockout of PU.1. Mice in which *Spi1* gene exon 5 was flanked by loxP sequences were bred with mice carrying a transgene containing Cre recombinase under the control of adiponectin (*Adipoq*) gene promoter and enhancer (Fig. 1A). Mice with adipose-specific knockout of PU.1 (adiponectinCre-PU.1^fl/fl^) and littermate control mice (PU.1^fl/fl^) were used for the study. To confirm ablation of aPU.1 expression, we isolated the adipocyte fraction from gonadal adipose tissue. As shown in Fig. 1B, PU.1 expression in the adipocytes of aPU.1KO mice was significantly down-regulated.

**Fig. 1.**
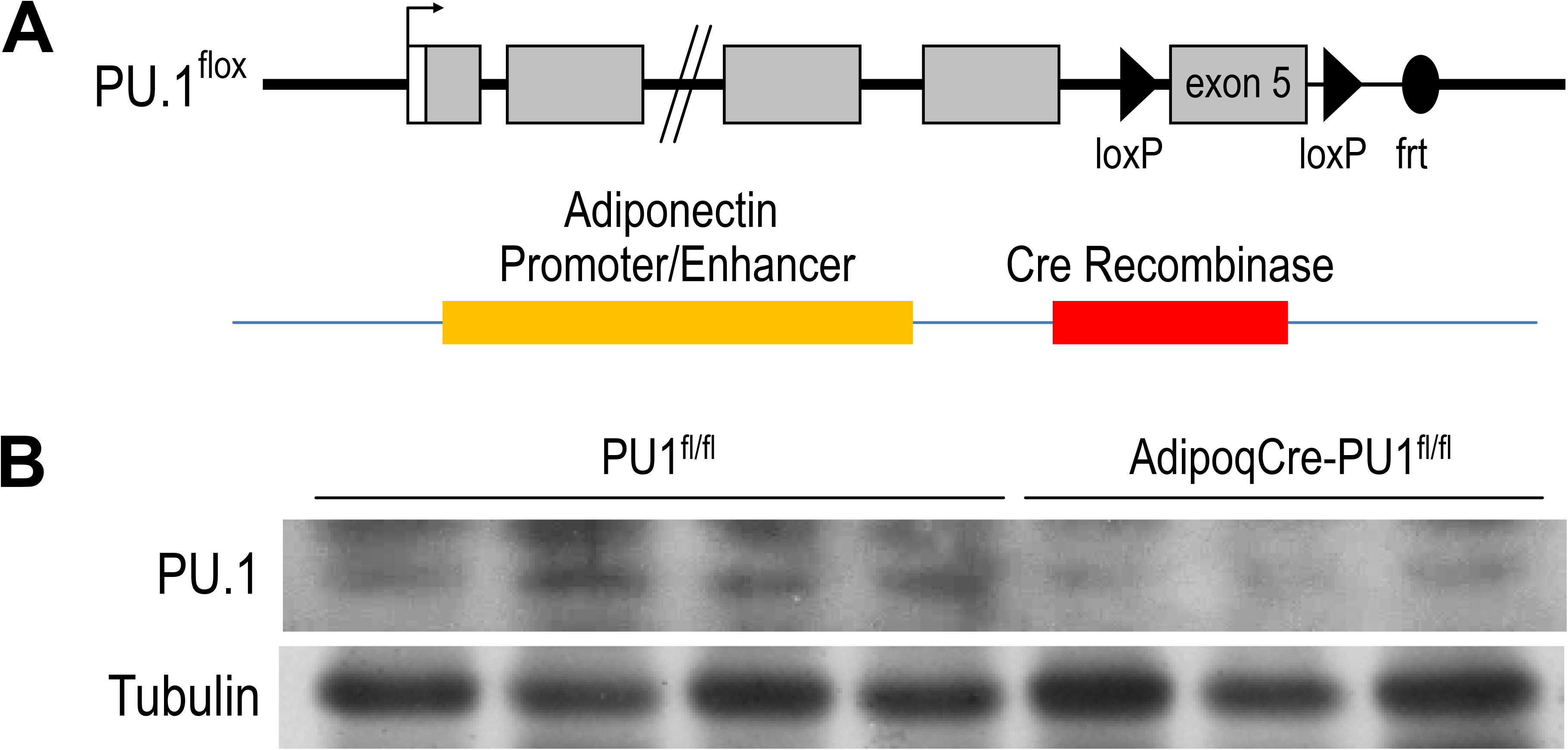
Generation of Adipocyte-Specific PU.1 KO Mice. (A) The schematic diagram of AdipoqCre-PU.1^fl/fl^ Mice. Mice that have the PU.1 gene exon 5 flanked with loxP sequences were breed with mice carrying a transgene containing Cre recombinase under the control of Adiponectin gene promoter and enhancer. (B) The expression of PU.1 in isolated adipocytes was detected using anti-PU.1 Western blot analysis.

### 3.2 Phenotype of young adult aPU.1KO mice

Young (4-5 mo) aPU.1KO male (Fig. 2A) or female (Supplementary Fig. 1A) mice exhibited no difference in body weight, lean body mass or fat mass from floxed littermate controls. Moreover, glucose tolerance tests found no difference in glucose homeostasis (Fig. 2B and Supplementary Fig. 1B) between 4-5 mo aPU.1 KO male or female mice and their floxed littermate controls. Using indirect calorimetry to measure energy expenditure however, we found that compared to wild-type littermates, male aPU.1KO mice had significantly higher average energy expenditure and basal energy expenditure under fasted and resting state (Fig. 2C). This increase in energy expenditure was not contributed by increased physical activity, as there was no difference in ambulatory activity in these mice (data not shown). In female aPU.1KO mice, no difference of energy expenditure was observed (Supplementary Fig. 1C).

**Fig. 2.**
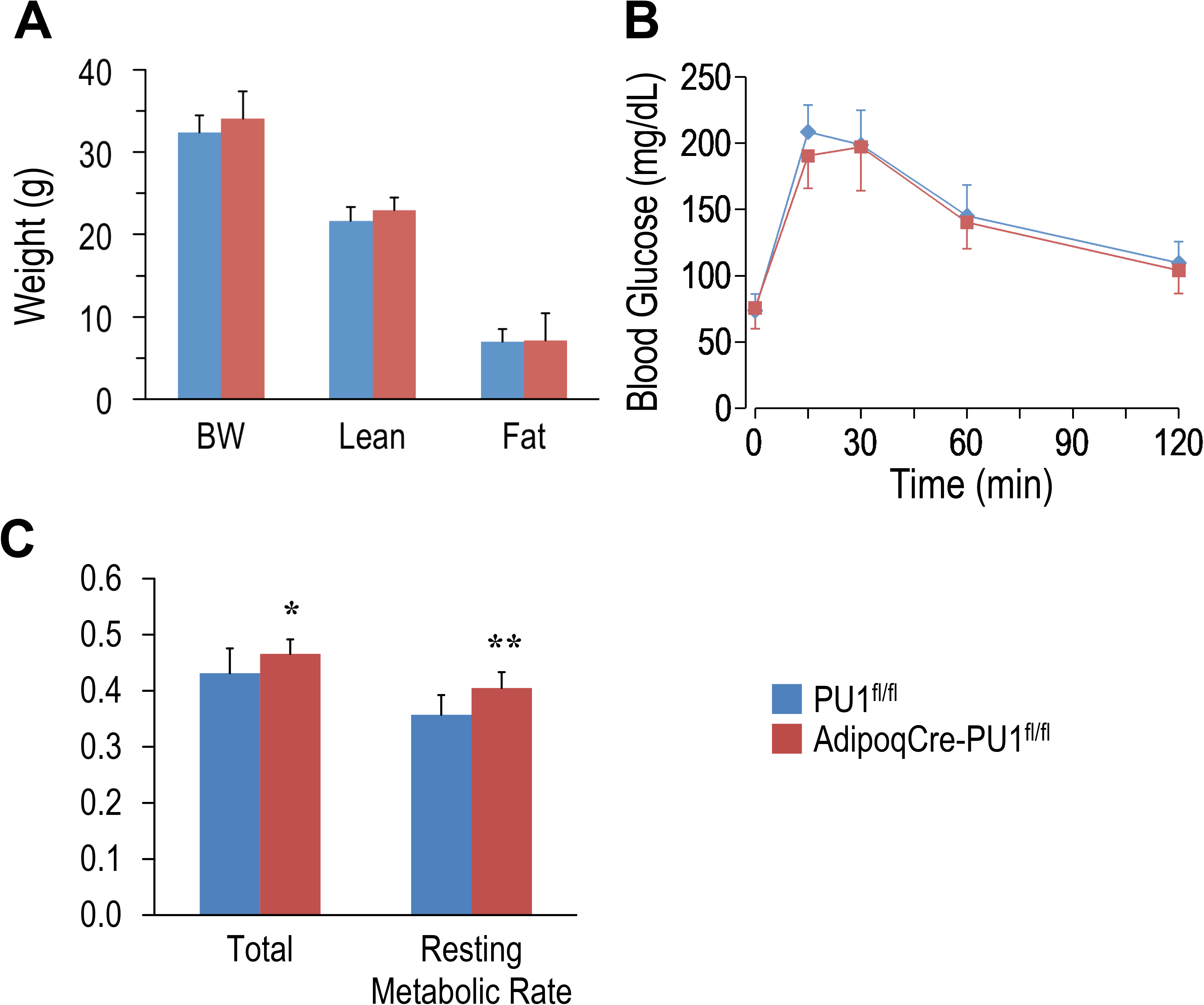
Adipocyte PU.1 Deficiency Has No Effect on Body Weight, Body Composition and Glucose Tolerance in Young Adult Male Mice. (A) Body weight and body composition of young adult (4-5 months of age) male AdipoqCre-PU.1^fl/fl^ mice and the control littermate PU.1^fl/fl^ mice. (B) For glucose tolerance test, mice were fasted overnight and injected with glucose. Blood glucose was then measured. (C) Energy expenditure was determined using indirect calorimetry in the integrated Comprehensive Lab Animal Monitoring System (CLAMS; Columbus Instruments) in 4-5 months old male AdipoqCre-PU.1^fl/fl^ mice and the control littermate PU.1^fl/fl^ mice. * p<0.05, ** p<0.01.

### 3.3 Deficiency of adipocyte PU.1 protects against age-associated obesity and glucose intolerance

At 10-11 months of age, control male mice gained significantly more body weight than aPU.1KO mice (Fig. 3A). Moreover, whereas lean body mass of aPU.1KO mice was indistinguishable from that of wild-type mice (Fig. 3A), fat mass was significantly lower (Fig. 3A), indicating that the difference of body weight was attributable primarily to a loss of adiposity. Fasted 1 yo aPU.1KO male mice exhibited significantly improved glucose tolerance (Fig. 3B) and insulin sensitivity (Fig. 3C) compared with WT controls. Female 10 mo aPU.1KO mice were similar to WT controls with respect to body weight and fat mass (Supplementary Fig. 2A) and insulin sensitivity (Supplementary Fig. 2C), although they did exhibit a modest improvement in glucose tolerance (Supplementary Fig. 2B).

**Fig. 3.**
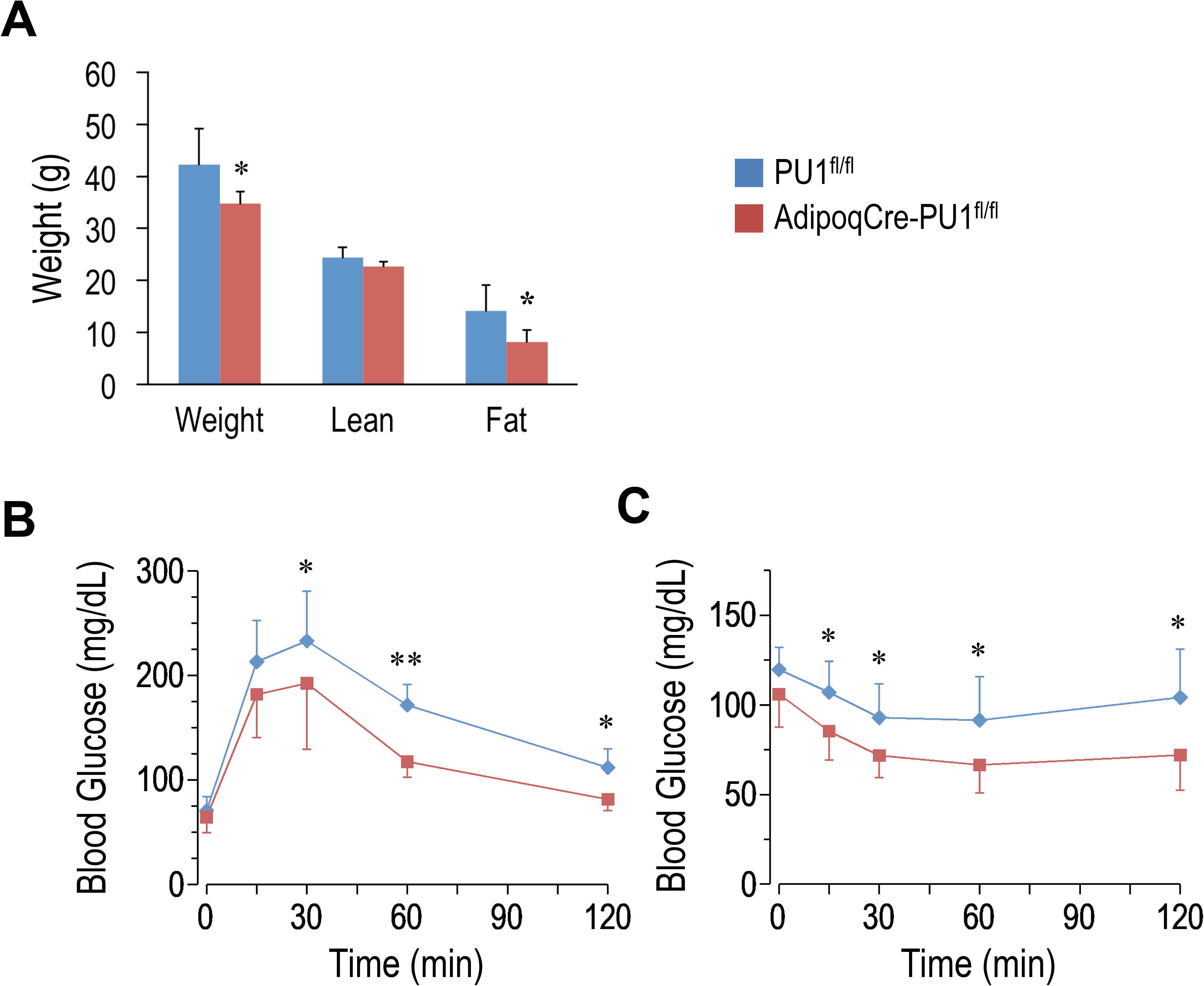
Adipocyte PU.1 Deficiency Protects Mice Against Age-Associated Obesity and Inuslin Resistance. (A) Body weight and body composition of one year old male AdipoqCre-PU.1^fl/fl^ mice and the control littermate PU.1^fl/fl^ mice. (B) For glucose tolerance test, one year old male mice were fasted overnight and injected with glucose (2g/kg body weight). Blood glucose was then measured. (C) For insulin tolerance test, mice were fasted for 4-hr and injected with insulin (0.8IU/kg body weight). Blood glucose was measured afterwards. * p<0.05, ** p<0.01.

### 3.4 Transcriptomic analysis identifies regulation of diverse metabolic transcriptional programs by PU.1

To investigate the metabolic phenotypes arising from loss of adipocyte PU.1, we performed RNA-sequencing of adipocytes isolated from epididymal adipose tissue of aPU.1KO and control mice. Significantly (*p*<0.01) induced and repressed genes were visualized on a volcano plot (Fig. 4A) and analyzed by Reactome Pathway Analysis (RPA; Fig. 4B). RPA analysis of down-regulated genes displayed an enrichment of pathways involved in extracellular matrix and immune signaling, including interleukin-10 signaling (Fig. 4B). Consistent with this, genes with well documented roles in these processes (*Il1rn*, *Il10ra*, *Il1b*, *Ccr1*, *Ccl6*) were highly repressed in aPU.1KO adipocytes (Fig. 4A). In contrast, up-regulated genes were enriched for pathways involved in adipogenesis and lipogenesis, including *Cebpa*, *Srebf1*/SREBP, *Nr1h3*/LXRα, *Mlxipl*/ChREBP, *Acly*, *Apoc1* and *Pck1* (Fig. 4A and Fig. 4B). These results were consistent with our previous findings of the roles of PU.1 in driving expression of inflammatory genes and transcriptional suppression of adipogenesis in cultured adipocytes ^19 20^.

**Fig. 4.**
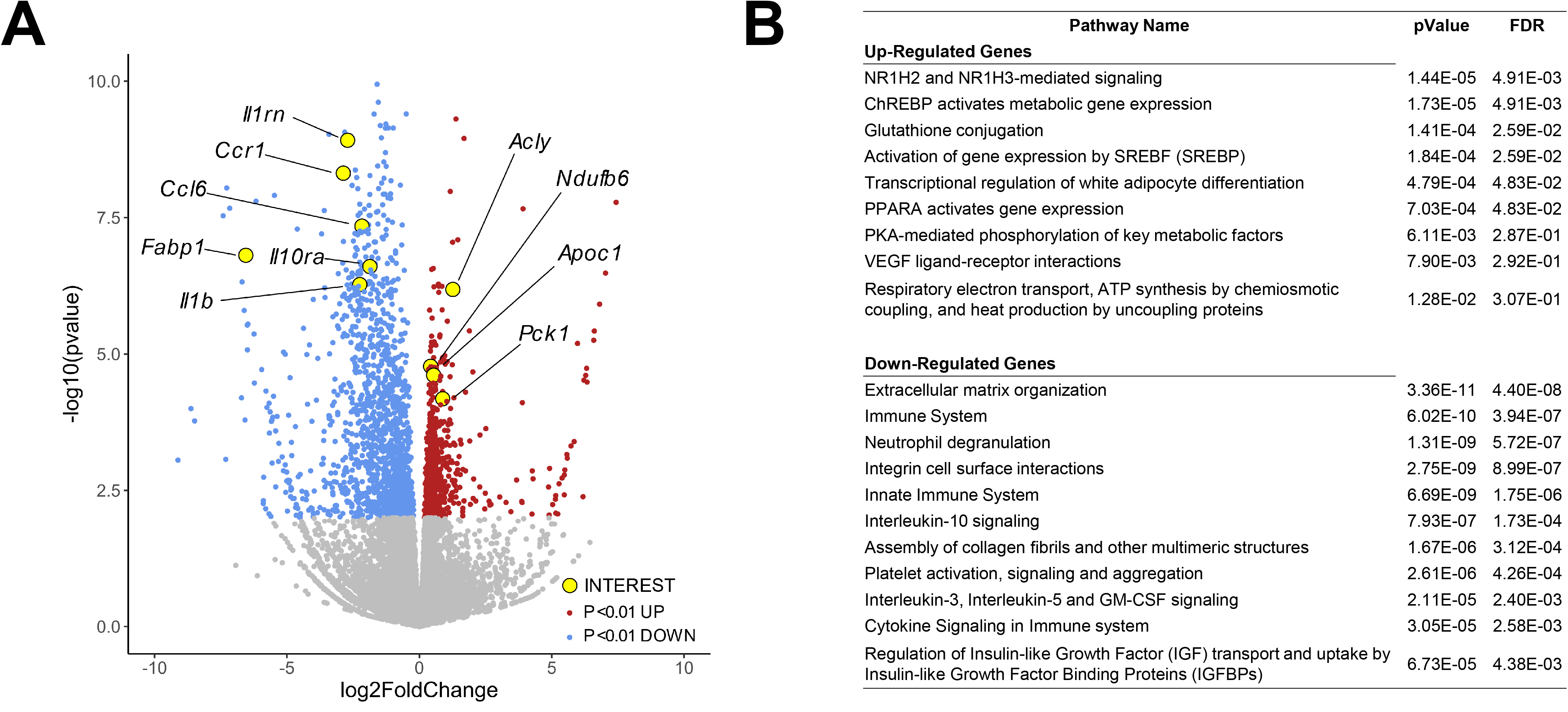
RNA-sequencing Analysis of Gene Expression Changes in the aPU.1KO Mice. (A) Volcano plot showing gene expression changes in adipocytes isolated from the gonadal adipose tissue, with key genes of interest highlighted in yellow. (B) Gene ontology analysis of differentially expressed genes (p<0.01) showing highly enriched pathways. The false discovery rate (FDR) is used for multiple hypothesis testing, with a standard cutoff of FDR<0.05 for significant pathway enrichment.

### 3.5 Transcriptional regulatory network analysis illuminates crosstalk of PU.1 with adipogenic and inflammatory signaling node networks

Although RPA analysis highlighted functional processes downstream of aPU.1-regulated genes (Fig. 4), it was less informative as to which were direct PU.1 transcriptional targets, and as to what aPU.1-modulated signaling pathways might regulate expression of these transcriptional targets. Signaling Pathways Project (SPP) consensomes are ranked consensus transcriptional signatures for signaling pathway nodes—receptors, enzymes, transcription factors and other nodes—computed from publicly archived ‘omics datasets ^27^. As such, consensomes have value in identifying potential high confidence transcriptional targets (HCTs) for specific nodes or node families in a given biological system^29^. To gain insight into members of the PU.1 transcriptional regulatory network, we next applied HCT intersection analysis to compute intersections between aPU.1KO-induced and aPU.1KO-repressed gene sets (FC >±1.5, *P*< 0.05) and a library of over 700 mouse HCT gene sets derived from archived transcriptomic or ChIP-Seq consensomes as previously described ^27, 29^. We interpreted the size and significance of these intersections as evidence for loss or gain of function of a given signaling node or node family in aPU.1KO adipocytes and, by inference, a functional relationship with aPU.1.

Figure 5A shows a heatmap displaying selected intersections between the aPU.1KO up and down gene sets and SPP mouse node family transcriptomic (upper table panel) and node ChIP-Seq (lower table panel) consensome HCTs. To assist in identifying candidate aPU.1-interacting nodes in WAT, Column P in Supplementary Table 2 represents percentiles of the mean WAT expression levels of each node derived from our RNA-Seq dataset. As an objective validation of our analysis, we benchmarked it against a list of 16 proteins identified by BioGRID ^31^ as mouse PU.1 interacting proteins (Supplementary Table 2, column Q). Figure 5B shows a regulatory footprint plot, in which signaling nodes that have significant HCT footprints with aPU.1KO-repressed genes are indicated in orange outline. In this plot, nodes that have the most highly enriched and significant regulatory footprints are located towards the top right of the plot. Reflecting the reliability of our predictions, the 16 BioGRID-sourced PU.1-interacting nodes (yellow data points in Fig. 5B) were enriched (OR = 11, *P*=2.6E-04) among nodes with significant (*Q*<1E-12) intersections with aPU.1-KO repressed genes (Fig. 5B). In addition to the expected prominent footprint for PU.1 itself in the down-regulated gene set (*Q*=1.9E-36), intersections with numerous canonical functional partners of PU.1 further validated our analysis. For example, robust intersections of aPU.1KO-repressed gene sets with transcriptomic HCTs for members of the toll-like ^32^, leptin ^33^, and chemokine ^34^ receptor and Protein kinase B/Akt ^35^ families are consistent with previous studies implicating signaling through these nodes in PU.1 function (Fig. 5A). Similarly, the increased energy expenditure of the aPU.1KO mice is reflected in the strong footprint within the aPU.1KO-induced gene set of transcriptomic HCTs for members of the PPARG coactivator 1 (PGC-1) family, which are well known mediators of metabolic control ^36^. Moreover, reflecting adrenergic stimulation of thermogenesis ^37^ as well as repression of inflammatory cytokines ^38^, we observed β-adrenergic footprints in both aPU.1KO-induced (*Q*=1E-4) and -repressed (*Q*=7E-33) gene sets (Fig. 5A).

**Fig. 5.**
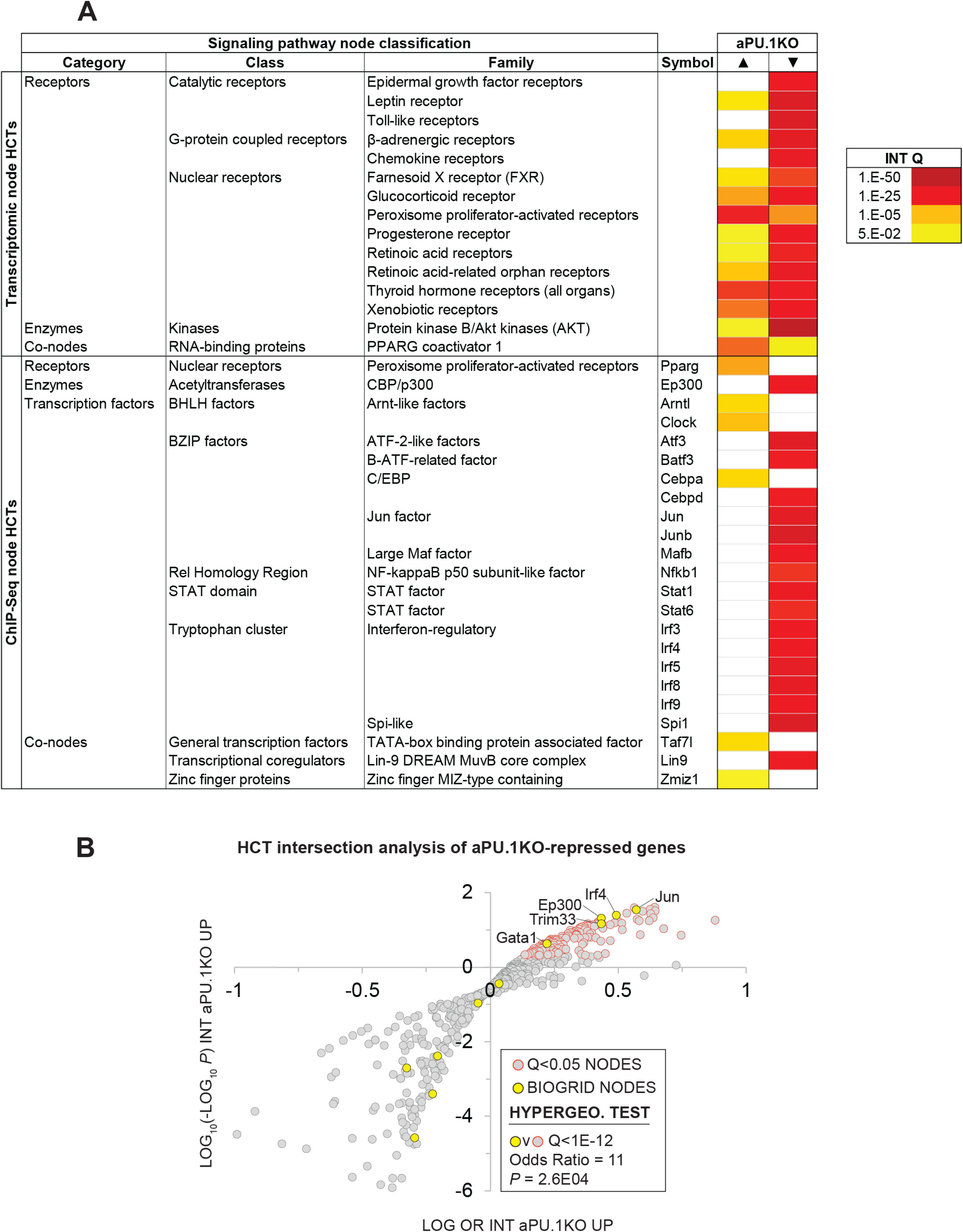
High Confidence Transcriptional Target (HCT) Intersection Analysis Identifies PU.1-Modulated Metabolic and Inflammatory Transcriptional Regulatory Hubs. HCT intersection Q-values (INT Q) for selected signaling pathway nodes or node families are indicated in the form of a heatmap. HCT intersection analysis was carried out as described in the Methods section. White cells represent Q > 5E-2 intersections. Full numerical data are in Supplementary Table 2.

Similarly, in the ChIP-Seq HCTs, robust intersections of aPU.1KO-regulated genes with HCTs for members of the interferon regulatory factor (IRF)^39^, STAT^40^, AP-1^41^, Atf^42^, C/EBP and GATA ^19^ transcription factor families are consistent with canonical PU.1 biology. Given our previous report that PU.1 functions synergistically with GATA transcription factors to inhibit adipogenesis ^19^, we were also interested to note intersections of aPU.1KO-regulated genes with HCTs for GATA family members (Fig. 5B). Congruent with our metabolic studies of aPU.1KO mice (Figs. 2 & 3), aPU.1KO-induced genes contained footprints for several nodes with familiar roles in the context of whole body energy metabolism, including Pparg (3E-7), Nr3c1/GR (1.2E-05) and Cebpa (1.2E-5). Strikingly, four nodes with known roles in circadian rhythms (Clock, 1.2E-5; Nr3c1/GR, 1.2E-05; Nr1d1/REV-ERB, 1.1E-04 and Arntl/BMAL1, 2.1E-04) also had appreciable intersections with the aPU.1KO-induced genes (Fig. 5A and Supplementary Table 2).

In addition to corroborating canonical PU.1 biology, our analysis suggested the possibility of crosstalk of PU.1 with signaling nodes with which it has no previously established functional relationships. For aPU.1KO-induced genes these included transcriptomic HCT intersections with the PGC-1 family and ChIP-Seq HCT intersections for Zbtb11, Taf3 and Zfp57. Similarly, for aPU.1KO-repressed genes, we noted intersections with HCTs for members of the E2A-related factor family and the MuvB complex members Lin9 and E2f4 (Supplementary Table 2).Of note, although some of the intersections were attributable to transcriptional induction or repression of their encoding genes in the absence of PU.1 (aPU.1KO v WT log2 FCs: *Cebpa*, 0.35; *Irf5*, -2.6; *Irf8*, -0.83; *Mafb*, -1.2; *Mef2c*, -0.95; Supplementary Table 1), the vast majority of nodes that had significant HCT intersections with aPU.1KO-regulated genes were not transcriptionally regulated. Collectively these data indicate repressive, protein-level cross-talk of PU.1 with adipogenic/lipogenic signaling nodes on the one hand, and on the other, positive cross-talk with distinct classes and families of inflammatory signaling nodes.

### 3.6 Consensome analysis identifies broad scale direct regulation by PU.1 of adipogenic gene expression

We previously showed that PU.1 inhibits adipogenesis^19^. With that in mind, our RNA-Seq analysis highlighted numerous aPU.1 KO-regulated genes that represented potentially novel, previously uncharacterized modulators of adipogenesis. We next wished to adopt a reduced-bias approach to explore this possibility in more detail. Using our previously-described consensome algorithm ^27^, we used five archived datasets (GSE2192, GSE60745, GSE14004, GSE12929, GSE20696) to generate a 3T3-L1 adipogenesis consensome, which ranks ∼12,500 genes according to the frequency with which they are upregulated or downregulated across independent transcriptomic 3T3-L1 adipogenic datasets (Supplementary Table 3 contains 3T3-L1 adipogenesis consensome genes with *P*<0.05, *n*=9152). Within the 3T3-adipogenesis consensome we designated Q<0.05 genes with a mean FC>2 (log FC > 1) as 3T3-L1 adipogenesis induced CTs (3T3-ADIPICTs, *n* = 508; Supplementary Table 3, column I) and Q<0.05 genes with a mean FC<0.5 (log FC < -1) as adipogenesis repressed CTs (3T3-ADIPRCTs, *n* = 100; Supplementary Table 3, column J). We first benchmarked the 3T3-L1 adipogenesis consensome against a set of 200 genes designated as hallmark adipogenesis-induced transcripts by the GSEA ^43^ resource (GSEA HALLMARK; Supplementary Table 3, column Q). Validating the 3T3-L1 adipogenesis consensome, the GSEA HALLMARK gene set was robustly over-represented among 3T3-ADIPICTs (OR = 10.1, *P* = 4.2E-58; Fig. 6A). Many of the aPU.1KO-regulated 3T3-ADIPICTs are immediately familiar in the context of adipogenesis, including *Cidec*^44^, *Adipoq* ^45^ and *Acsl1*^46^. Interestingly, numerous aPU.1KO-induced (Supplementary Table 3, column R) and aPU.1KO-repressed (Supplementary Table 3, column S) genes have 3T3-L1 adipogenesis consensome rankings that are comparable to or exceed those of classic adipogenic markers, but have potential roles in adipogenesis that to date are unexplored in the research literature (Table 1).

**Fig. 6.**
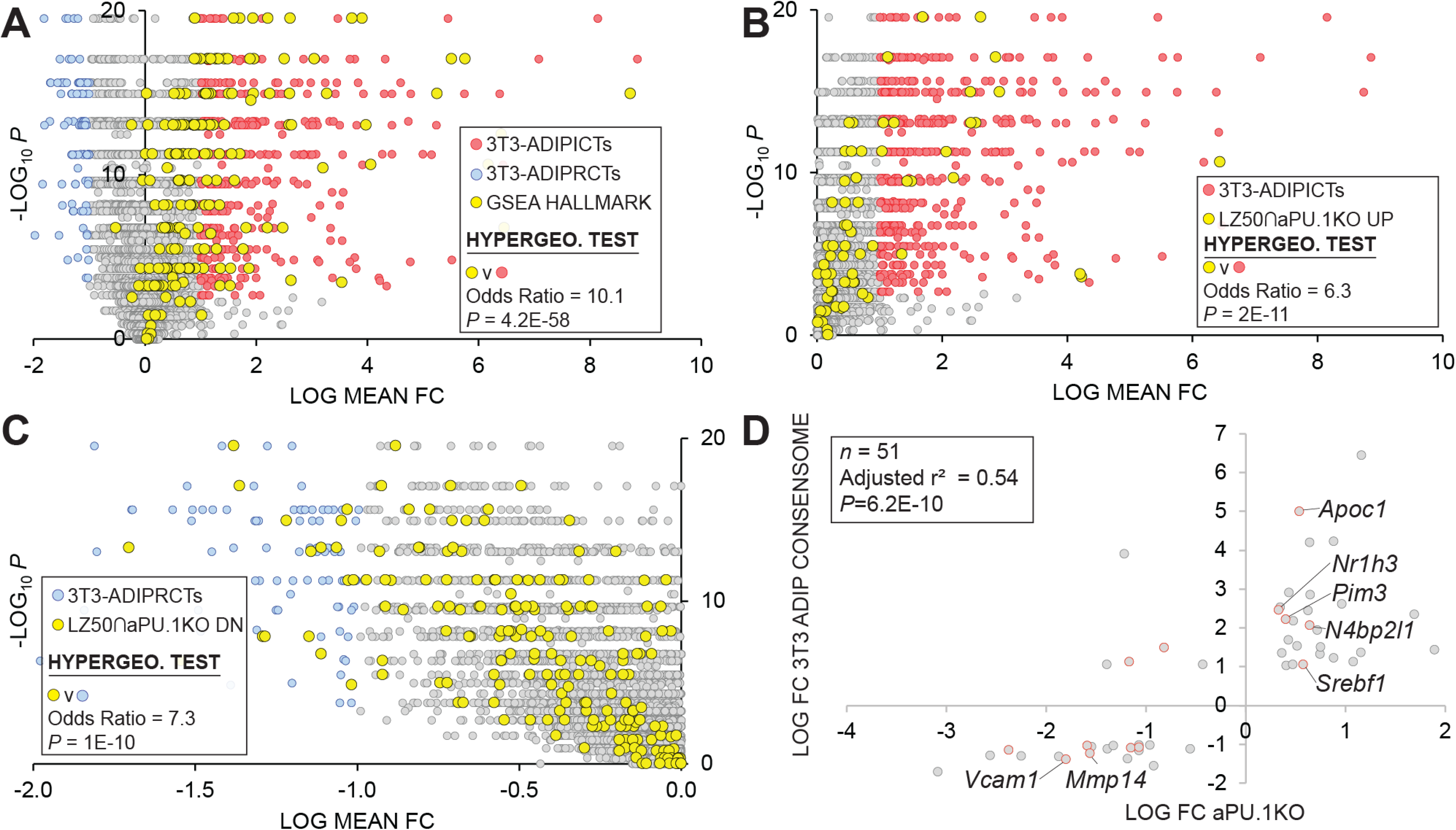
3T3-L1 Adipogenic Differentiation Consensome. The mouse 3T3-L1 adipogenesis transcriptomic consensome ranks mouse genes based on their discovery rates across five independent, publicly archived 3T3-L1 adipogenesis transcriptomic datasets. Hypergeometric test odds ratio and associated *P*-value are indicated in each panel. A. Validation of the 3T3-L1 adipogenesis consensome against the GSEA adipogenesis Hallmark gene set. B. Over-representation of aPU.1KO-induced LZ50 genes among 3T3-ADIPICTs. C. Over-representation of aPU.1KO-repressed LZ50 genes among 3T3-ADIPRCTs. D. Correlation between aPU.1KO log FC and adipogenesis log mean FC.

**Table 1.**

| Target |  |  |  | aPU.1KO | 3T3-ADIP CONSENSOME |  |  |
| --- | --- | --- | --- | --- | --- | --- | --- |
| Category | Class | Family | Symbol | log2 FC | log2 FC | %ile | Q-VALUE |
| aPU.1KO-induced |  |  |  |  |  |  |  |
| Receptors | G protein coupled | G protein-coupled receptor | Gpr176 | 0.45 | 1.27 | 99 | 6.3E-15 |
| Enzymes | Dehydratases | 3-hydroxyacyl-CoA dehydratases | Hacd4 | 0.40 | 1.04 | 92 | 5.6E-10 |
| Co-nodes | Apoptosis | Niban apoptosis regulator | Niban1 | 0.64 | 2.07 | 90 | 5.6E-10 |
|  | Developmental proteins | Testis development related protein | Tdrp | 0.45 | 0.89 | 87 | 1.7E-08 |
|  | Membrane proteins | Tetraspanin | Tspan12 | 0.48 | 2.18 | 87 | 1.7E-08 |
|  | Other co-nodes | MAM domain containing | Mamdc2 | 0.37 | 0.57 | 94 | 1.4E-11 |
|  | Stress response factors | DnaJ heat shock protein (Hsp40) member | Dnaja3 | 0.40 | 0.67 | 99 | 6.3E-15 |
| aPU.1-repressed |  |  |  |  |  |  |  |
| Receptors | Ligands | FAT atypical cadherin | Fat4 | -1.13 | -1.01 | 90 | 5.6E-10 |
| Enzymes | ADP ribosyltransferases | Poly [ADP-ribose] polymerases (PARP) | Parp14 | -0.56 | -1.11 | 95 | 1.2E-11 |
|  | Dehydrogenases | 3 beta hydroxysteroid dehydrogenases | Hsd3b7 | -1.07 | -1.15 | 80 | 8.0E-07 |
|  | GTPases | RAB, member RAS oncogene | Rab7b | -2.56 | -1.28 | 80 | 8.0E-07 |
|  | Reductases | Glyoxylate and hydroxypyruvate reductases | Grhpr | -3.09 | -1.71 | 95 | 1.2E-11 |
|  | Regulatory factors | NEDD4 binding proteins like | N4bp2l1 | -2.25 | -1.29 | 80 | 8.0E-07 |
| Co-nodes | Cytoskeleton | FERM domain containing | Frmd4a | -0.92 | -1.54 | 74 | 1.8E-05 |
|  | Glycoproteins | Glycoprotein integral membrane | Ginm1 | -1.07 | -1.06 | 95 | 1.2E-11 |
|  | Membrane proteins | CKLF like MARVEL transmembrane domain | Cmtm3 | -1.55 | -1.05 | 97 | 3.3E-13 |
|  | Pleckstrin domain | Pleckstrin homology domain containing | Plekho2 | -1.59 | -1.03 | 98 | 1.1E-13 |
|  | Transporters | Vesicle amine transport | Vat1 | -0.86 | -1.11 | 80 | 8.0E-07 |

To focus on direct aPU.1 targets among genes with elevated rankings in the 3T3-adipogenesis consensome, we next percentilized peak call heights (*n*=679) from a previously published PU.1 3T3-L1 ChIP-Seq analysis by Lazar and colleagues ^47^ and mapped these to genes in the 3T3-L1 adipogenesis consensome (Supplementary Table 3, column T). For the purposes of subsequent statistical analyses, the 50^th^ percentile of this gene set (*n* = 444) is referred to here as LZ50. Consistent with broad, direct antagonism by PU.1 of the 3T3-L1 adipogenic transcriptional program, we observed robust enrichment of aPU.1KO-induced LZ50 genes (*n*=78) in 3T3-ADIPICTs (Fig. 6B; OR=6.3, *P*=2.2E-11) and of aPU.1KO-repressed LZ50 genes (*n*=293) among 3T3-ADIPRCTs (Fig. 6C; OR=7.3, *P*=1E-10). Further reflecting the role of aPU.1 as an important direct regulator of adipogenic differentiation, we observed a clear positive correlation (adjusted r^2^ = 0.54, *P*=6E-10) between aPU.1KO log FCs and mean 3T3-adipogenesis consensome log FCs for 3T3-ADIPRCTs or 3T3-ADIPICTs in the 50^th^ percentile of PU.1 3T3-L1 peaks (*n* = 51. Fig 6D). Studying these 51 genes further, we identified two prominently pro-adipogenic members of the 3T3-ADIPICTs subset, *Srebf1* (encoding SREBP1) and *Nr1h3* (encoding LXRα), that have not been previously appreciated as direct PU.1 transcriptional targets. Similarly, uncharacterized candidate direct PU.1 targets within the 3T3-ADIPRCT gene subset included *Vcam1*, previously shown to mediate adhesion of inflammatory macrophages to adipocytes as a potential mechanism driving insulin resistance ^48^, and *Mmp14*, whose role in adipogenic collagen turnover has been linked to obesity ^49^. Collectively, our *in vivo* analysis confirms and adds value to previous in vitro studies implicating PU.1 as an anti-adipogenic, pro-inflammatory driver of gene expression in adipocytes.

### 3.7 SPP web resource facilitates the generation of novel hypotheses around PU.1 regulation of adipogenic gene expression

To make full use of the adipogenesis consensome for hypothesis generation around aPU.1 regulation of adipogenic gene expression, it is important that the data points be placed in the context of data points from other mouse adipose transcriptomic experiments. For each gene in Supplementary Table 3 therefore, column U links to an SPP website interface showing the specific experimental data points for that gene from the 3T3-L1 adipogenesis expression profiling datasets. In addition, the interface includes data points from transcriptomic experiments in mouse adipose tissue or cell lines involving genetic or small molecule perturbation of various receptor and enzyme signaling nodes, as well as data points from metabolic challenges such as cold exposure. For insight into nodes directly regulating expression of genes in the adipogenesis consensome, Supplementary Table 3 column V links to an interface showing data points from ChIP-Seq experiments carried out in mouse adipose tissue or cell lines. Data points in both the transcriptomic and ChIP-Seq interfaces link to contextual pop-up windows, which in turn point to the full source datasets on the SPP website.

### 3.8 Confirmation of PU.1 regulation of pro-inflammatory cytokines and thermogenesis genes in adipose tissues

Given the global transcriptional repression of cytokine production in aPU.1KO adipocytes indicated by the RNA-Seq analysis (Fig. 4), we next used quantitative real-time RT-PCR (Q-PCR) to validate this observation. We confirmed the reduction of the expression of *Ccl2/*MCP-1 and *Tnf* in adipocytes isolated from epididymal WAT of 1 yo male aPU.1KO mice (Fig. 7A). Since aPU.1KO mice exhibited elevated energy expenditure, we speculated that we might also observe induction of thermogenic expression programs in brown adipose tissue (BAT) of aPU.1KO mice. Consistent with this hypothesis, Q-PCR analysis identified transcriptional induction in aPU.1KO BAT of two key thermogenic genes, *Ppargc1a*/PGC-1α and *Ucp1* (Fig. 7B). These results suggest a molecular mechanism underlying PU.1 regulation of adipocyte inflammation and thermogenesis, and offer an explanation of the phenotype of the aPU.1KO mice.

**Fig. 7.**
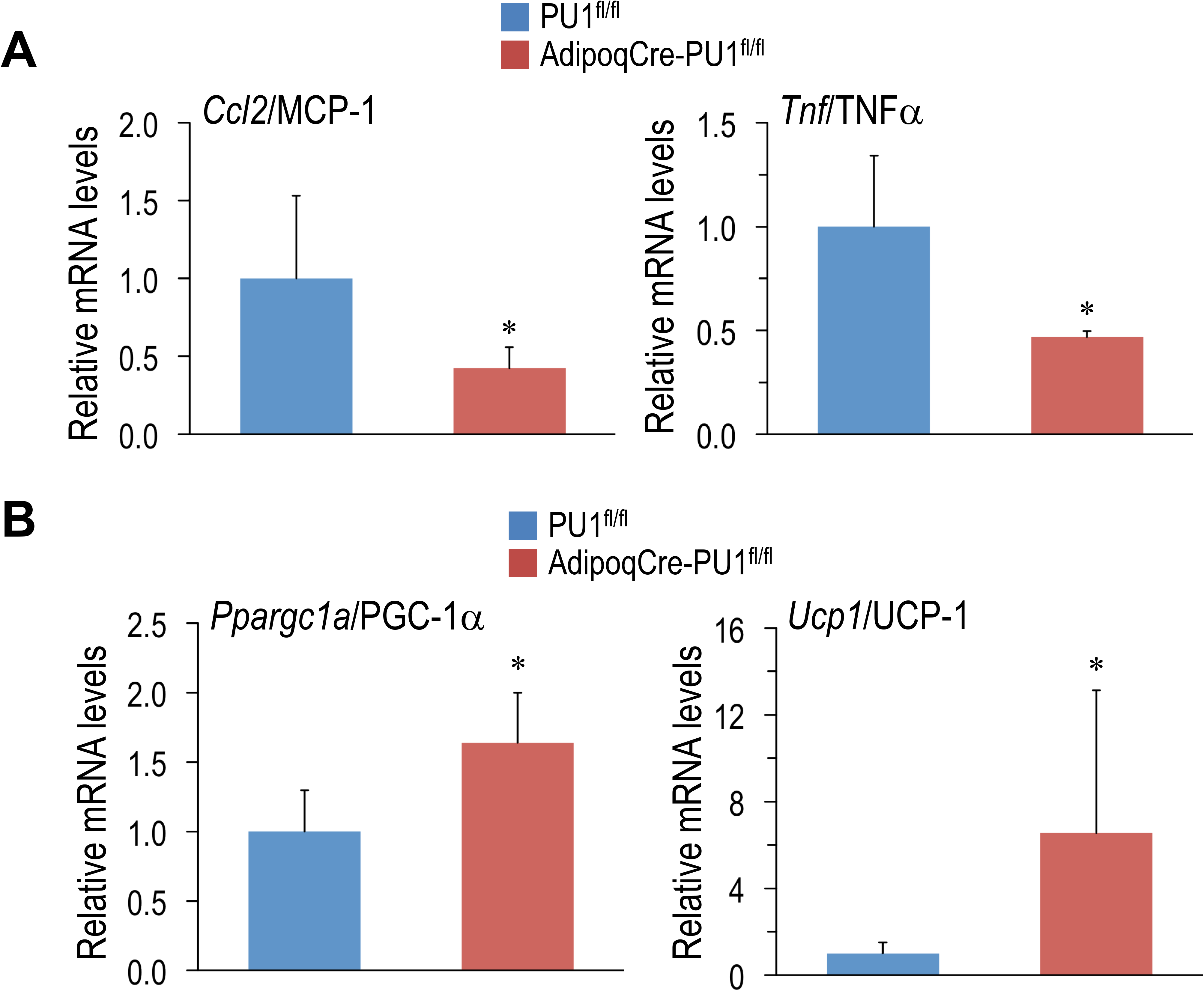
Loss of PU.1 in Adipocytes Reduces Inflammatory Gene Expression and Promotes Thermogenic Expression. (A) Adipocytes were isolated from the gonadal adipose tissue and mRNA expression of *ccl2*/MCP-1 and TNFα were measured using real-time RT-PCR. (B) Brown adipose tissue mRNA expression of PGC-1α and UCP1 and TNFa were measured using real-time RT-PCR. *p<0.05

### 3.9 Integrated analysis of aPU.1KO transcriptional regulatory networks and mouse phenotypes

The Mammalian Phenotype Ontology (MPO) ^50^, uses evidence from the research literature to assign specific metabolic and physiological functions to the products of mouse genes. As such, MPO annotations represent a potentially powerful approach to inferring the metabolic impact of transcriptional regulatory networks in the aPU.1KO mouse. Given the increased energy expenditure of aPU.1KO mice (Figure 2), we first wished to gather evidence for nodes that are transcriptional mediators of increased whole body energy expenditure in these mice. To do this we examined the intersection between nodes that had significant intersections with aPU.1KO-UP genes, and those encoded by genes whose disruption in mice mapped to the MPO term “abnormal energy expenditure” (AEE; Supplementary Table 2, column R). Consistent with the increased energy expenditure in the aPU.1KO mice, nodes encoded by AEE-mapped genes were strongly over-represented among nodes that have *Q*<0.01 intersections with aPU.1KO-induced genes (OR=10.6, *P*=4E-5, Fig. 8A). These nodes included the nuclear receptors Pparg^51^ and Ppara^52^, the circadian clock regulators Clock^53^ and Arntl^54^, and the C/EBP family member Cebpa^55^ which, as previously noted, was strongly transcriptionally induced in adipocytes in the aPU.1KO mice (Fig. 2).

**Fig. 8.**
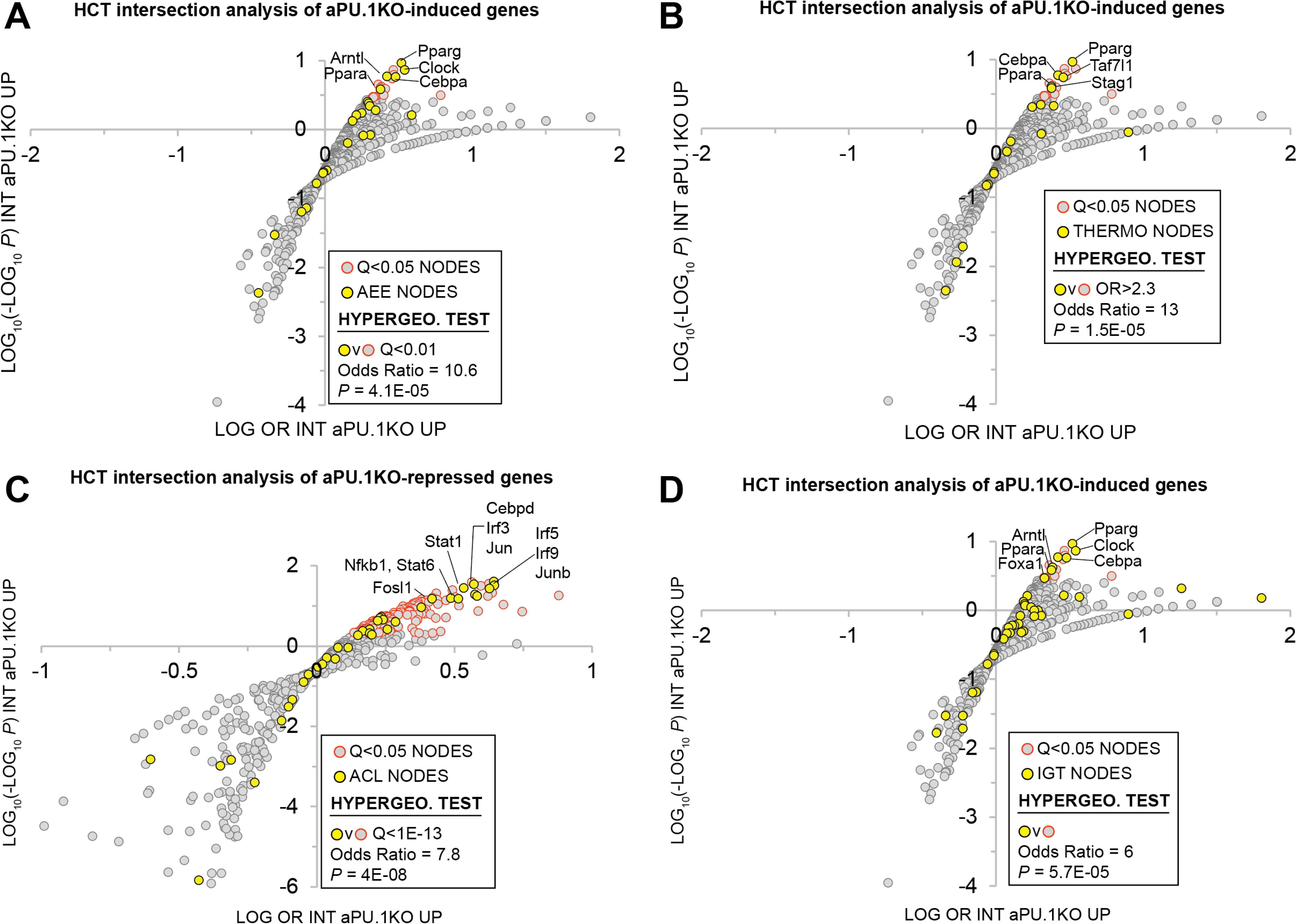
Mammalian Phenotype Ontology Analysis Of Nodes With Significant HCT Intersections With aPU.1KO-Induced and Repressed Gene Sets. A. Enrichment of nodes encoded by genes that map to the MPO term “abnormal energy expenditure” (AEE) among nodes that have significant HCT intersections with aPU.1KO-induced genes. B. Enrichment of nodes encoded by genes that map to the MPO terms “decreased brown adipose tissue amount”, “decreased core body temperature” or “impaired adaptive thermogenesis” (collectively referred to as THERMO genes) among nodes that have significant HCT intersections with aPU.1KO-induced genes. C. Enrichment of nodes encoded by genes that map to the MPO term “abnormal cytokine levels” (ACL) among nodes that have significant HCT intersections with aPU.1KO-repressed genes. D. Enrichment of nodes encoded by genes that map to the MPO term “impaired glucose tolerance” among nodes that have significant HCT intersections with aPU.1KO-induced genes. Please refer to the Methods for details of the analysis.

Next, to identify potential transcriptional mediators of thermogenic gene expression in the aPU.1KO mice (Fig. 4), we compared nodes that had significant intersections in aPU.1KO-UP genes with those encoded by a set of genes that mapped to at least one of the MPO terms “decreased brown adipose tissue amount”, “decreased core body temperature” or “impaired adaptive thermogenesis” (collectively referred to as THERMO genes; Supplementary Table 2, column S). Consistent with the induction of BAT thermogenic genes in the aPU.1KO animals, nodes encoded by THERMO genes were strongly over-represented among those that have *Q*<0.01 intersections with aPU.1KO-induced genes (OR=13, *P*=1.5E-05; Fig. 8B). These included the nuclear receptor Pparg^56^, the C/EBP family transcription factor Cebpa^55^, the general transcription factor Taf7l ^57^ and the cohesin complex member Stag1 ^58^. Collectively these data indicate that a non-redundant role for PU.1 in repressing a transcriptional regulatory network driving thermogenic gene expression *in vivo*.

Node HCT intersection analysis had previously indicated that numerous inflammatory transcription factors with roles in cytokine production were functionally impacted by the loss of PU.1 (Fig. 5A). To gain insight into the PU.1 adipocyte transcriptional hub supporting inflammatory cytokine production *in vivo*, we compared nodes that had significant intersections with aPU.1KO-DOWN genes and those encoded by genes whose disruption in mice mapped to the MPO term “abnormal cytokine levels” (ACL; Supplementary Table 2, column T). Reflecting transcriptional repression of numerous inflammatory cytokine genes in the aPU.1KO mice (Fig. 4, Fig. 7A, and Supplementary Table 1), nodes encoded by ACL genes were strongly over-represented among those with Q<1E-13 intersections with aPU.1KO-repressed genes (OR=7.85, *P*=4E-08; Fig. 8C). These included members of the IRF (Irf3, Irf5), STAT (Stat1, Stat6) and AP-1 (Jun, Junb) transcription factor families, as well as Fosl1, Cebpd and Nfkb1. Collectively these data indicate that PU.1 has a non-redundant role in anchoring a transcriptional regulatory network supporting adipocyte cytokine gene expression in vivo.

We showed that aPU.1KO mice exhibited improved glucose homeostasis (Fig.3). To gather evidence for the specific signaling nodes contributing to this phenotype, we compared nodes that had significant intersections in aPU.1KO-UP genes with those encoded by genes mapped to the MPO term “impaired glucose tolerance” (IGT; Supplementary Table 2, column U). Consistent with improved glucose homeostasis in the aPU.1KO mice, nodes encoded by IGT genes were strongly over-represented among nodes with Q<0.01 intersections with aPU.1KO-induced genes (OR=6, *P*=5.7E-05, Fig. 8D). Confirming the link between improved glucose tolerance and energy expenditure, these included the five previously identified AEE nodes (Pparg, Ppara, Clock, Arntl and Cebpa), in addition to Foxa1.

## 4. DISCUSSION

In this study, we set out to investigate the role of the transcription factor PU.1 in adipocytes *in vivo,* particularly during aging. We observed that although young (4-5 mo) aPU.1KO mice had no overt phenotypes with respect to body weight, body composition, or glucose homeostasis, they did exhibit elevated energy expenditure. Moreover, at around one year of age, aPU.1KO mice were protected against age-associated obesity, adipose tissue inflammation, and insulin resistance. Mechanistically, and consistent with their elevated energy expenditure, we found that loss of adipocyte PU.1 suppressed inflammatory transcriptional programs in WAT and promoted thermogenic gene expression in BAT. Using a combination of conventional, literature-based pathway analysis and a novel ‘omics dataset-centric analytic platform, we identified numerous PU.1-modulated signaling systems and downstream functional pathways that shed mechanistic light on these phenotypes.

Consistent with the enhanced energy expenditure of aPU.1KO mice, we identified a robust transcriptional footprint within aPU.1KO-induced genes for members of the PGC-1 family (Fig. 5A), which are well known transcriptional drivers of thermogenesis and energy metabolism ^59^. This represented strong evidence, validated by subsequent Q-PCR analysis (Fig. 7A), that PU.1 suppresses thermogenic transcriptional programs in mice. To afford insight into the functional pathways involved, Supplementary Table 2 column L indicates aPU.1KO-induced genes that are HCTs for the PGC-1 family. Many of these are familiar players in cellular energy metabolism that have emerged from studies in the research literature. The induction of *Ndufb6*, *Ndufb10* and *Ndufs6*, for example, reflects our finding from RPA analysis that respiratory electron transport chain pathway genes were enriched among aPU.1KO-induced genes (Fig. 4A). Moreover, upregulation of the transferrin (*Trf*) gene may reflect potential endocrine or paracrine signaling from the white adipocytes to activate brown or beige adipocyte thermogenesis in aPU.1KO mice ^60^. The power of consensome analysis, however, is that it illuminates genes for which a specific function is not described in the research literature, but for which, based upon close regulatory relationship with a node family computed from ‘omics datasets, that function can be inferred with a high level of confidence. This is the case with genes such as *Blcap*, *Gkap1*, *Proca1* and others, for which, as aPU.1KO-induced PGC-1 HCTs, a functional contribution to enhanced bioenergetics of the aPU.1KO animals can be reasonably surmised. Supporting this assertion, and confirming the clinical relevance of our study, the human ortholog of *Fam13a* (aPU.1KO logFC = 1.3; PGC-1 family consensome 95^th^ percentile) has been recently shown to regulate fat distribution and metabolic traits through its action on adipose tissue ^61^. In summary, we conclude that activation of PGC-1 family signaling in aPU.1KO mice contributes to the increased energy expenditure, reduced adiposity, improved glucose homeostasis and insulin sensitivity of aPU.1KO mice, and ultimately protects against age-related metabolic abnormalities in these animals.

Transcriptomic analysis has cast PU.1 in the global maintenance of pro-inflammatory transcriptional programs in response to sepsis (14) or lipopolysaccharide stimulation (15). Although transcriptional induction by PU.1 of pro-inflammatory factors, including *Tnf* (27), *Il1b* (28, 29) and *Il6* (30), as well as *Ccl3* (31) and *Ccl2* (24) has been extensively documented in macrophages and other immune cell lineages, there is increasing interest in its role as a pro-inflammatory transcription factor in adipose tissue. Building on our previous study showing that PU.1 activates inflammatory cytokine expression in cultured adipocytes ^20^, our current study shows that depletion of adipocyte PU.1 results in broad, transcriptome-scale suppression of inflammatory programs in adipocytes. Since inflammation is a key mediator for insulin resistance and metabolic syndrome ^1^, depletion of this inflammatory transcriptional program likely contributes to the phenotypes of the aPU.1KO mice. Going to the underlying mechanism, we found that the expression levels of many proinflammatory cytokines driven by the PU.1 transcription factor, such as *il1b*, *tnf* and *ccl2*/MCP-1, are down regulated in the adipocytes of aPU.1KO mice (Fig. 4 and Fig.8). IL-10 signaling, which has been shown to inhibit thermogenesis and energy expenditure in adipocytes ^62^, was also downregulated in PU.1-deficient adipocytes (Fig. 4B). Members of the NOD-like family of receptors have prominent roles in the transcriptional regulation of inflammasome pathways. Our transcriptomic analysis identified significant down-regulation of genes encoding two members of the NOD-like family, *Ciita* and *Nrpl3,* in aPU.1KO adipocytes (Supplementary Table 1). Similarly, consistent with the strong TLR regulatory footprint in the aPU.1KO-repressed genes (Fig. 5A), genes encoding four members of the TLR family (*Tlr1*, *Tlr6*, *Tlr7* and *Tlr8*) are repressed in the aPU.1KO. Given that TLRs ^63^, *Ciita* ^64^ and *Nrpl3* ^65^ are known to be induced in obese adipose tissue or to support adipocyte inflammation, it can be justifiably speculated that their transcriptional induction in adipocytes makes an important contribution to PU.1’s action in promoting inflammation and insulin resistance.

On a broader scale, HCT intersection analysis (Fig. 5A), validated by integration with literature-based mouse phenotype annotations (Fig. 8C), reflects the profound impact of loss of PU.1 on the function of numerous inflammatory node families. For example, aPU.1KO-downregulated genes contain a sizeable regulatory footprint for members of the IRF transcription factor family (Fig. 5A and Supplementary Table 2). Given that the roles of members of the IRF family in the regulation of adipogenesis, inflammation and thermogenesis in adipocytes are well-documented ^66, 67^, we interpret the presence of this footprint as evidence for strong, network-level interactions between PU.1 and IRFs in adipocytes. On the other hand, transcription factors that suppress inflammatory gene expression, such as PPARγ (16, 17) and LXRs (18, 19), have DNA binding sites adjacent to PU.1 binding sites, potentially reflecting mutual functional antagonism. Given that we observed evidence for activation of both PPAR and LXR in response to aPU.1 loss of function (Fig. 4B), we speculate that aPU.1 may also drive inflammation through the suppression of PPAR and LXR transcription factors.

Although ‘omics datasets have intrinsic value for metabolic research, they realize their full value when integrated with existing data resources to facilitate the generation of hypotheses around metabolic signaling pathways not explored in the research literature. A unique aspect of our RNA-Seq dataset is that rather than limiting it to a standalone analysis of aPU.1KO-regulated genes, we have placed it in the context of millions of regulatory data points curated from archived ‘omics datasets by the SPP cell signaling knowledgebase ^26^. Annotation of aPU.1KO-regulated gene list (Supplementary Table 1) according to the SPP classification, for example, provides for an immediate appreciation of the diversity of cellular functions impacted by PU.1 depletion. Similarly, HCT intersection analysis (Fig. 5 and Supplementary Table 2) affords a unique perspective on the various receptors, enzymes, transcription factors and co-nodes that are functionally impacted by PU.1 depletion and which, by extension, are candidate PU.1-interacting proteins. Finally, the adipose-centric SPP transcriptomic and ChIP-Seq Regulation Reports to which the 3T3-L1 adipogenic consensome (Supplementary Table 3) links provide the user with a rich, contextual perspective to generate hypotheses around transcriptional regulation of novel effectors of adipose tissue biology. By integrating these three data resources in a single study, we provide for a unique perspective on PU.1-dependent transcriptional regulatory networks in adipocytes, and an appreciation of how diverse signaling nodes impact expression of a specific PU.1 target gene ^2726^.

The collective value of our supplementary material to researchers in generating novel metabolic hypotheses can be illustrated with reference to *Gpr176*, identified in Supplementary Table 1 as a gene encoding a member of the G protein-coupled receptor family that is strongly transcriptionally dependent upon PU.1. With the exception of a role in the regulation of circadian clock in the suprachiasmatic nucleus ^68^, the function of this receptor is largely uncharacterized. The 3T3-L1 adipogenesis consensome ranks *Gpr176* 445^th^ of 12525 genes (mean log FC -1.71, CQV 1E-11, 95^th^ %ile), indicating that *Gpr176* is robustly and consistently downregulated during adipogenic differentiation. The SPP transcriptomic Regulation Report for *Gpr176* (Supplementary Table 3 column U) contains data points documenting its regulation by prominent regulators of lipid metabolism, including FGF21, PPARG and members of the PGC-1 family. Similarly, the ChIP-Seq Regulation Report (Supplementary Table 3 column V) provides evidence for direct regulation of *Gpr176* by PU.1 and numerous other nodes, including Polycomb group proteins and members of the C/EBP, STAT and BRD families. Finally, the enrichment among aPU.1KO-induced genes of HCTs for numerous characterized transcriptional regulators of circadian rhythms (Arntl/BML1, Nr1d1/Rev-Erbα, Clock, and Nr3c1/GR; Fig 5A and Supplementary Table 2) suggests that PU.1 regulation of circadian transcriptional programs in adipocytes may well extend beyond *Gpr176*. Set in the context of existing evidence documenting circadian connections between adipose tissue biology, lipid metabolism and the immune system ^69–71^, the SPP data points suggest a hypothesis implicating PU.1 as a transcriptional co-ordinator of circadian programs in adipocytes and immune cells. Indeed, such a notion is supported by a previous report of global enhancement of PU.1 transactivation in Arntl/BML1-depleted macrophages^72^.

The recent characterization of PU.1 as a transcriptional driver of fibrosis ^73^ is interesting given the known role of fibrosis in supporting the inflammatory state ^74^ and obesity ^75^. Interestingly, RPA identified a strong repression in the aPU.1KO adipocytes of pathways related to the extracellular matrix (ECM), a critical player in the development of fibrosis ^76^ (Fig. 4B). Inspecting the aPU.1KO-repressed genes more closely, we identified three members of the fibrinogen family (OR=318, *P*=1E-08, hypergeometric test), six members of the integrin family (OR=7.5; P=1E-04, hypergeometric test), 12 members of the cluster of differentiation group (OR = 13, *P*=1E-10, hypergeometric test) and nine members of the collagen family (OR=6.9, P=5E-06, hypergeometric test), many of which have been implicated in fibrosis and obesity ^77–85^. Most intriguingly of all perhaps, aPU.1KO-repressed genes contain eight members of the major urinary protein (MUP) family (OR=417, P=4E-21), several of which are among the most strongly repressed genes. MUP proteins are related to members of the lipocalin family, which have documented connections to a variety of fibrotic conditions ^86–88^, as well as obesity and insulin resistance ^89, 90^. Collectively, our analysis data point to a pivotal role for PU.1 in driving fibrotic transcriptional programs that support inflammatory pathways in adipocytes.

Our observation that PU.1 plays a role in the development of age-associated metabolic syndrome shed light on not only a novel PU.1 action in adipocytes, but also the nature of age-related metabolic defects. Metabolic changes developed during the aging process share similarities with that caused by obesity, but also possess some unique characteristics ^22^. The underlying mechanism is not well characterized. A recent study identified a sub-population of adipocytes present only in the subcutaneous adipose tissue of aged mice or humans ^91^. These cells have elevated PU.1 expression, which causes defective adipogenesis and proinflammatory cytokines secretion to inhibit adipogenesis of neighboring cells. This finding is in agreement with our results, supporting an important role of PU.1 in aging adipose tissue, in the development of age-associated adipocyte dysfunction, with a likely consequence of whole-body metabolic defects.

As an adipocyte-specific knockout, our model underscores the contribution of adipocyte-autonomous functions of PU.1 to disorders of systemic metabolism. However, PU.1 in tissues other than adipose may also contribute to metabolic syndrome. For example, expression of hepatic PU.1 is also elevated in diet-induced obese and diabetic mice, and is positively correlated with insulin resistance and liver inflammation in humans ^92^. Depletion of PU.1 in non-parenchymal liver cells, likely in liver macrophages, inhibited liver inflammation, hepatic steatosis and whole body insulin resistance ^92^. Taken together, our study identifies PU.1 as a regulator of metabolic syndrome in both adipocytes and in liver macrophages. Therefore, PU.1 is an important driver for metabolic disorders in adult and older animals and may serve as a therapeutic target for the treatment of metabolic syndrome.

## AUTHOR CONTRIBUTIONS

Conceptualization: TQ. Investigation: KC, ADA, XG, and WP. Validation: EL and DS. Methodology/Software: SAO and NM. Formal analysis: AM, SAO and NJM. Writing and editing: TQ, AM, YS, and NJM. Funding acquisition: TQ and NJM.

## ACKNOWLEDGMENTS

This work was supported by a US Department of Agriculture grant (3092-5-001-059), NIH (DK075978) and AHA (18TPA34170539) to Q. T., by NIH DK097748 to N.J.M., and by the DKNET Summer Of Data student internship, supported by DK097748, to A.M. These sponsor play no role in study design; in data acquisition, analysis and interpretation; in the writing of the manuscript; and in the decision to submit for publication.

## Supplemental Figures

**Suppl Fig. 1.**
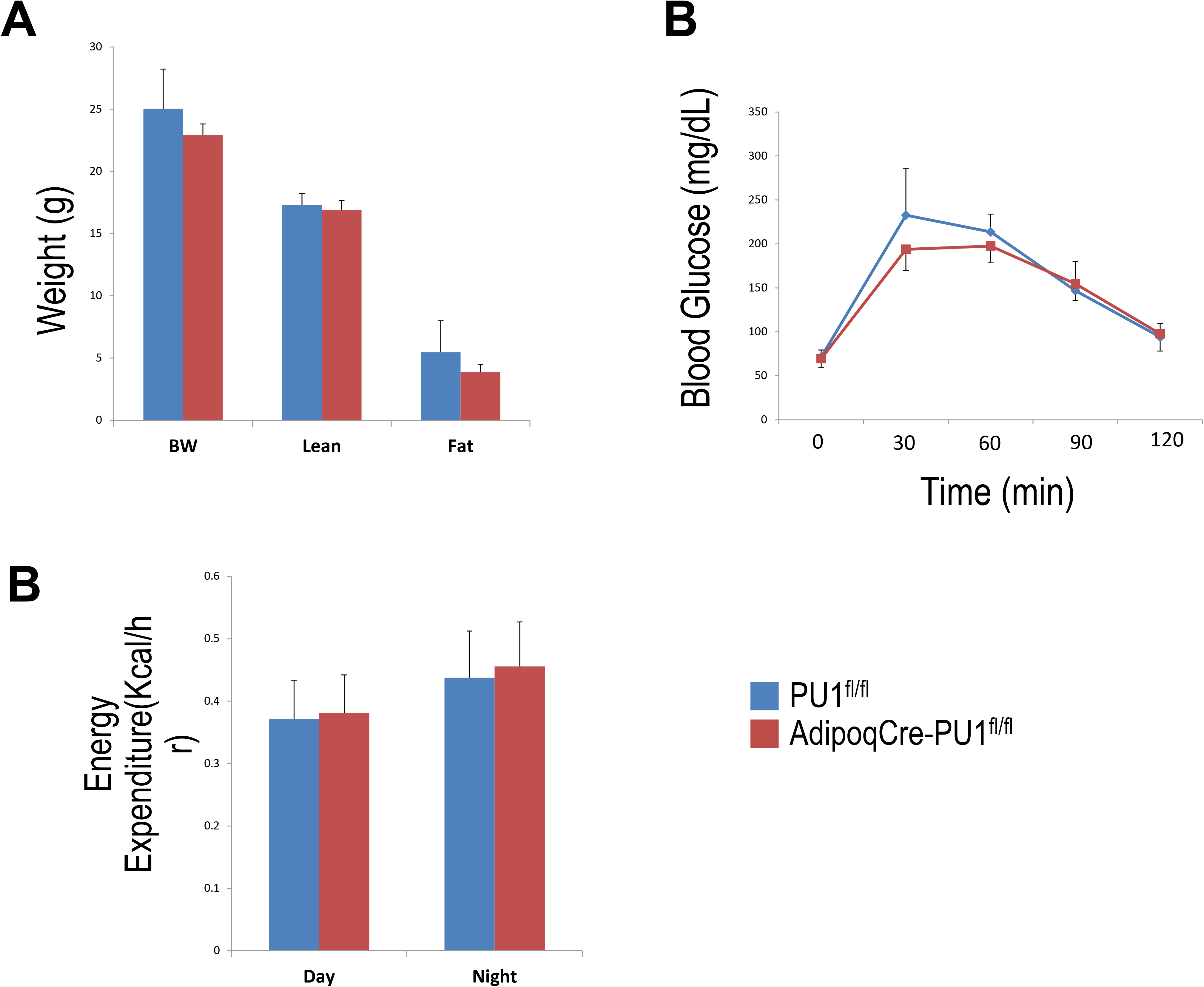
Adipocyte PU.1 Deficiency Has No Effect on Body Weight, Body Composition and Glucose Tolerance in Young Adult Female Mice. (A) Body weight and body composition of young adult (4-5 months of age) female AdipoqCre-PU.1^fl/fl^ mice and the control littermate PU.1^fl/fl^ mice. (B) For glucose tolerance test, mice were fasted overnight and injected with glucose. Blood glucose was then measured. (C) For insulin tolerance test, mice were fasted for 4 hrs and injected with insulin. Blood glucose was then measured. *P<0.05

**Suppl Fig. 2.**
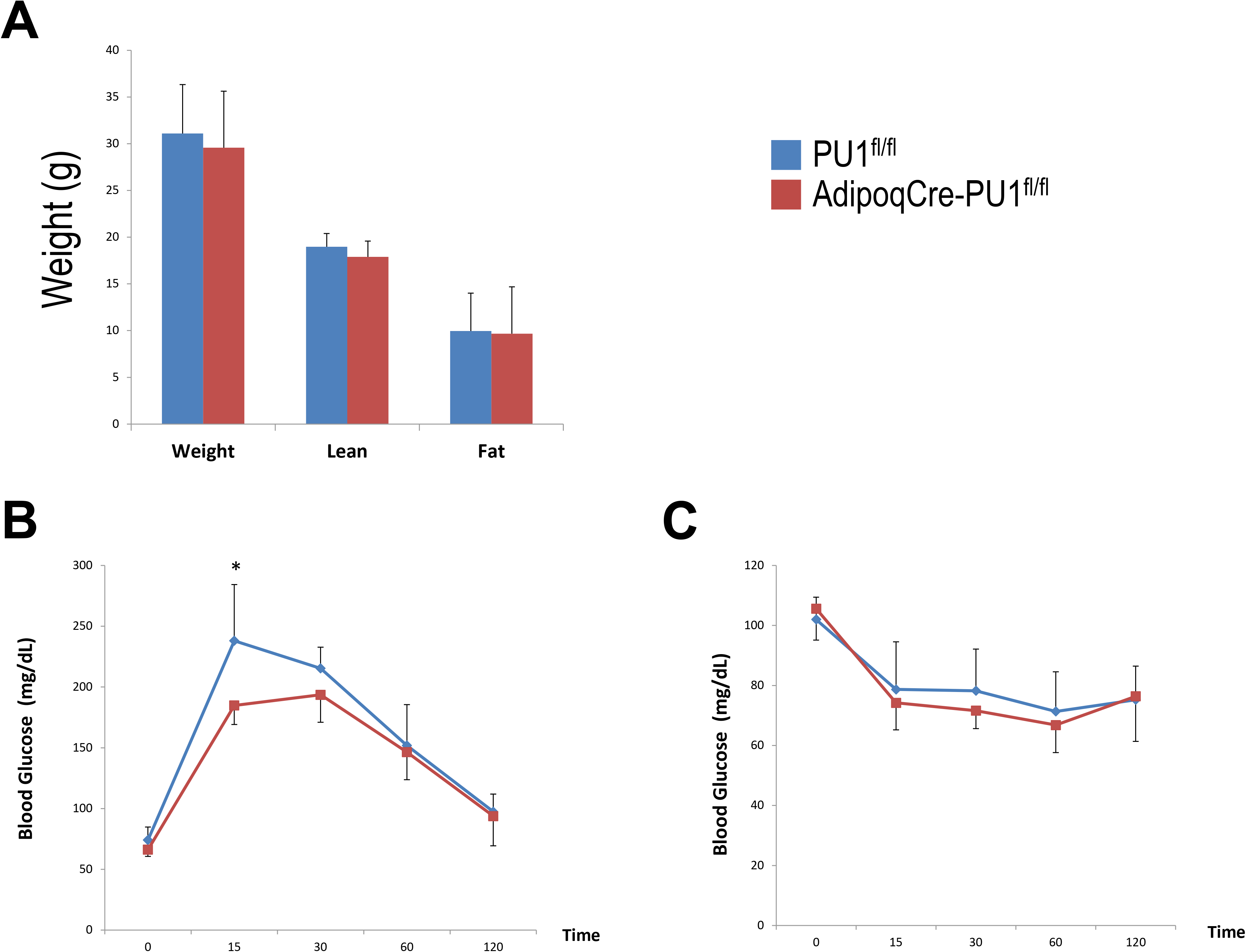
Adipocyte PU.1 Deficiency Has minimal Effect on Body Weight, Body Composition and Glucose Tolerance in Older Female Mice. (A) Body weight and body composition of older adult (10-11 months of age) female AdipoqCre-PU.1^fl/fl^ mice and the control littermate PU.1^fl/fl^ mice. (B) For glucose tolerance test, mice were fasted overnight and injected with glucose (2g/kg body weight). Blood glucose was then measured. (C) For insulin tolerance test, mice were fasted for 4 hrs and injected with insulin (1.0 IU/kg). Blood glucose was then measured. *P<0.05

